# Structures of Naked Mole-Rat, Tuco-Tuco, and Guinea Pig Ribosomes–Is rRNA Fragmentation Linked to Translational Fidelity?

**DOI:** 10.1101/2025.08.01.667930

**Authors:** Cristina Gutierrez-Vargas, Swastik De, Suvrajit Maji, Zheng Liu, Zhonghe Ke, Martina Nieß, Andrei Seluanov, Vera Gorbunova, Joachim Frank

**Author notes:** contributed equally (Cristina Gutierrez-Vargas, Swastik De).

## Abstract

Ribosomes are central to protein synthesis in all organisms. Among mammals, the ribosome functional core is highly conserved. Remarkably, two rodent species, the naked mole-rat (NMR) and tuco-tuco display fragmented 28S rRNA, coupled with high translational fidelity and long lifespan. The unusual ribosomal architecture in the NMR and tuco-tuco has been speculated to be linked to high translational fidelity. Here we show, by single-particle cryo-electron microscopy (cryo-EM), that despite the fragmentation of their rRNA, NMR and tuco-tuco ribosomes retain their core functional architecture. Compared to ribosomes of the guinea pig, a phylogenetically related rodent without 28S rRNA fragmentation, ribosomes of NMR and tuco-tuco exhibit poorly resolved, certain expansion segments. In contrast, the structure of the guinea pig ribosome shows high similarity to human ribosome. Enhanced translational fidelity in the NMR and tuco-tuco may stem from subtle, allosteric effects in dynamics, linked to rRNA fragmentation.

## Introduction

Ribosomes, as the central players in protein synthesis, have a highly conserved core across species, especially within the critical functional centers: the decoding site, the peptidyl transferase center, and tRNA-binding sites (Frank 2000; Doudna et al., 2002; Ramakrishnan 2002; Steitz 2008; Jobe et al., 2019). Outside of this conserved core, ribosomal composition can vary considerably across species, particularly in peripheral regions. In eukaryotes, such species-specific adaptations are largely due to unique insertions or extensions in ribosomal proteins (r-proteins) and distinct rRNA divergent domains or expansion segments (ESs) (Melnikov et al., 2012).

Eukaryotic rRNA genes are present in multiple tandem copies in the nucleus, with each unit containing the genes for 18S, 5.8S, and 28S rRNAs and their flanking transcribed spacers, which are removed during processing. Beyond the typical excision of spacers in standard rRNA processing, some organisms undergo additional, more unusual RNA cleavage events that result in fragmented rRNAs. rRNA fragmentation has been noted in organisms from diverse taxa— including bacteria, protozoa, and insects. Notably, the vertebrate lineage was largely assumed to lack such internal rRNA fragmentation in mature ribosomes (Fujiwara and Ishikawa, 1986). However, new findings have revealed a unique situation in the South American rodents, tuco-tuco, where the 28S rRNA undergoes processing through a mechanism akin to splicing, resulting in a single break within an intron located in the D6 domain (Melen et al., 1999).

Azpurua et al. (2013) subsequently reported an unexpected processing event in the 28S ribosomal RNA (rRNA) of the naked mole-rat (NMR) (*Heterocephalus glaber*), the longest-lived rodent with the maximum lifespan of 41 years (Buffenstein & Ruby, 2021). The NMR 28S rRNA is uniquely cleaved into two fragments, approximately 2.5 and 3 kb in size, resulting from two cuts within the D6 region (also known as the Divergent Regions or Expansion Segments) of the rRNA gene. This processing excises a 263-nt fragment containing a unique 118-nt sequence. The authors suggested this excised sequence may have originated from a transposable element, supported by flanking direct repeats. While the exact mechanism of cleavage remains unknown, possibilities include ribozyme activity or a specific ribonuclease. This form of fragmented rRNA is primarily observed in lower eukaryotes and prokaryotes, with the tuco-tuco (*Ctenomys talarum*), another rodent, being the only other known vertebrate example (Burgin et al., 1990; Rawson et al., 1971; Stevens and Pachler, 1972; Eckert et al., 1978; Castro et al., 1981; Spencer et al., 1987; van Keulen et al., 1991; Zarlenga and Dame, 1992; Applebaum et al., 1966; Ishikawa and Newburgh, 1972; Shine and Dalgarno, 1973; Lava–Sanchez and Puppo, 1975; Jordan et al., 1976; Ware et al., 1985; Boer and Gray, 1988; Fujiwara and Ishikawa, 1986; Azpurua et al., 2013).

Translational fidelity refers to the accuracy of amino acid incorporation into a polypeptide chain, according to the codon sequence encoded in mRNA. Thus, fidelity has two components: one is the error rate of aminoacyl-tRNA (aa-tRNA) synthetases, or their ability to accurately match amino acids to their corresponding tRNAs, and the other is the ribosome’s ability to accurately match tRNA anticodons to mRNA codons (Liu et al., 2017). Azpurua et al. (2013) specifically measured the second component, as the luciferase assay used assessed misincorporation at sense codons, premature stop codon readthrough, and frameshifting, all of which are ribosome-mediated. Azpurua et al. (2013) suggested that the NMR’s higher translational fidelity could be responsible for its longevity. This aligns with the “error catastrophe theory of aging” (Orgel, 1963; Edelmann and Gallant, 1977), which posits that the accumulation of errors in protein synthesis can eventually impair an organism’s function. We hypothesized that the unique 28S rRNA cleavage in the NMR might enhance translational fidelity by affecting ribosome dynamics. We also speculated that high translation fidelity might indirectly contribute to the NMR’s long lifespan and disease resistance by reducing protein misfolding (Buffenstein et al., 2002).

Here we set out to examine the ribosomal structures of the NMR and tuco-tuco. We have also included a phylogenetically related rodent, the guinea pig (*Cavia porcellus*), that does not have the cleaved 28S RNA as a control. NMRs, apart from their exceptional translational fidelity, which they share with tuco-tucos (Ke et al., 2017), are renowned for their unusual biological features, including extreme longevity, cancer resistance, and a remarkable tolerance of hypoxic conditions (Buffenstein 2005; Azpurua et al., 2013; Liang et al., 2010; Seluanov, et al., 2009). These traits make NMRs a valuable model for studying the molecular mechanisms of aging and disease resistance. Tuco-tucos, with their diverse species groups and high chromosomal variability, have adapted to subterranean environments in a way that provides unique insights into the processes of evolution and speciation (Prada et al., 2011; Torgasheva et. al., 2017; MacManes et al., 2012). Guinea pigs, meanwhile, have long been an essential model organism in biomedical research, owing to their immune system’s close resemblance to that of humans (Padilla-Carlin et al., 2008; Morrison et al., 2018). Together, these rodents offer a wealth of physiological and evolutionary diversity, allowing comparative studies that advance our understanding of cellular function and potential links to human health and disease.

To date, no high-resolution ribosome structures have been reported for the NMR, tuco-tuco, and guinea pig. Our study fills this important knowledge gap, enriching the range of mammalian ribosome structures available for evolutionary studies. Understanding diversity among ribosomal structures in model organisms is essential for molecular medicine, particularly in drug development. Ribosomes are largely conserved, but even small structural differences can influence how drugs interact with the translation machinery. Variations among species could lead to differential drug responses, potentially introducing inaccuracies into preclinical studies.

Examining ribosomal differences helps refine drug development strategies, improving predictions for human treatment outcomes and enhancing the overall accuracy of preclinical research.

Our cryo-EM structures of NMR and tuco-tuco ribosomes display differences from the guinea pig ribosome structure such as in mobile, poorly visible, certain expansion segments. These differences may be linked to enhanced translation fidelity, however, current data does not provide a direct link. It is likely, although not directly observable by cryo-EM, that fragmentation may change the dynamics of the ribosome during aa-tRNA selection and accommodation such that these steps occur with improved accuracy.

## Results

### 1. Sequence curation of naked mole-rat, tuco-tuco, and guinea pig ribosomal rRNAs

To investigate evolutionary divergence in the 28S ribosomal RNA (rRNA) across rodent species, we reconstructed the 28S rRNA sequences for NMR and tuco-tuco using experimental and publicly available data. For the NMR, we previously generated partial 28S rRNA sequence data using Rapid Amplification of cDNA Ends (RACE) from brain-derived RNA (Azpurua et al., 2013). For tuco-tuco, we extracted total RNA and submitted it for RNA sequencing (RNA-seq) at the JP Sulzberger Columbia Genome Center. Using the resulting RNA-seq data, we recovered nearly the entire 28S rRNA transcript. In contrast, full-length annotated 28S rRNA sequences for mouse and guinea pig were retrieved from the RNAcentral database.

Sequence alignment revealed that our partial NMR 28S rRNA corresponds to nucleotides 1711– 2600 of the mouse 28S rRNA (based on mouse numbering, Supplementary Figure X) (Azpurua et al., 2013). The extracted tuco-tuco sequence spans nearly the entirety of this region, missing only a few small segments at the 3′ end.

Pairwise percent identity analysis was performed across two regions flanking an NMR-specific insertion (positions 1934–1990, based on mouse numbering). In the upstream region (1711– 1933), tuco-tuco exhibited greater sequence similarity to guinea pig (94.74%) and mouse (85.96%) than to NMR (83.33%). NMR shared only 80.70% identity with guinea pig and mouse both, in this region. In the downstream region (1991–2582), following the insertion, a marked drop in similarity between NMR and the other species was observed. For example, NMR and mouse showed only 77.30% identity, whereas mouse and guinea pig maintained a high similarity of 88.54%.

The NMR-specific insertion (positions 1934–1990), which is absent in mouse, tuco-tuco, and guinea pig, was excluded from this apparent divergence analysis (i.e. based on % identity). Interestingly, even outside of this insertion, NMR sequences exhibit reduced similarity to other species. These results suggest that the NMR 28S rRNA has undergone accelerated evolutionary changes, diverging substantially from both phylogenetically close and distant rodent species. Notably, mouse and guinea pig remain the most similar among the species studied, with tuco-tuco consistently intermediate between them and NMR.

### 2. Overall structures of the naked mole-rat, tuco-tuco, and guinea pig ribosomes

In our structural analysis of the NMR, tuco-tuco, and guinea pig 80S ribosomes, we identified distinct conformational states in each species, differentiated by tRNA occupancy, and presence and degree of intersubunit rotation. For the NMR ribosome, three major states emerged: (i) a non-rotated ribosome with E-site tRNA (nrt80S), (ii) a rotated (13.2° for the head and 9.7° for the body) ribosome with P/E-site tRNA and eEF2 (rt80S-P/E-eEF2), and (iii) a rotated (4.4° for head, 8.0° for body) ribosome with P/E tRNA alone (rt80S-P/E) where A/P-tRNA density is weak, likely indicating heterogeneity. Similarly, the tuco-tuco ribosome displays three comparable conformational states, with variations in both intersubunit rotation (13.7° for head, 7.4° for the body, and 1.7° for head and 10.2° for body, respectively) and tRNA occupancy, as well as weak A/P tRNA density in the rotated state. In contrast, two primary conformations are observed for the guinea pig ribosome: (i) non-rotated state with E-site tRNA and fragmented density for P-site tRNA, and (ii) a slightly rotated (2.5° for head, 9.6° for body) state with E-site tRNA only. The conformational states of the NMR ribosome were resolved at resolutions of 3.1 Å, 3.4 Å, and 3.5 Å, while the tuco-tuco ribosome achieved resolutions of 3.0 Å for the non-rotated state, and 3.1 Å for both rotated states. The guinea pig ribosome was similarly resolved (<3.7 Å) to high structural detail, enabling comprehensive modeling of ribosomal proteins and rRNA regions across all three species (Figure 1).

**Figure 1:**
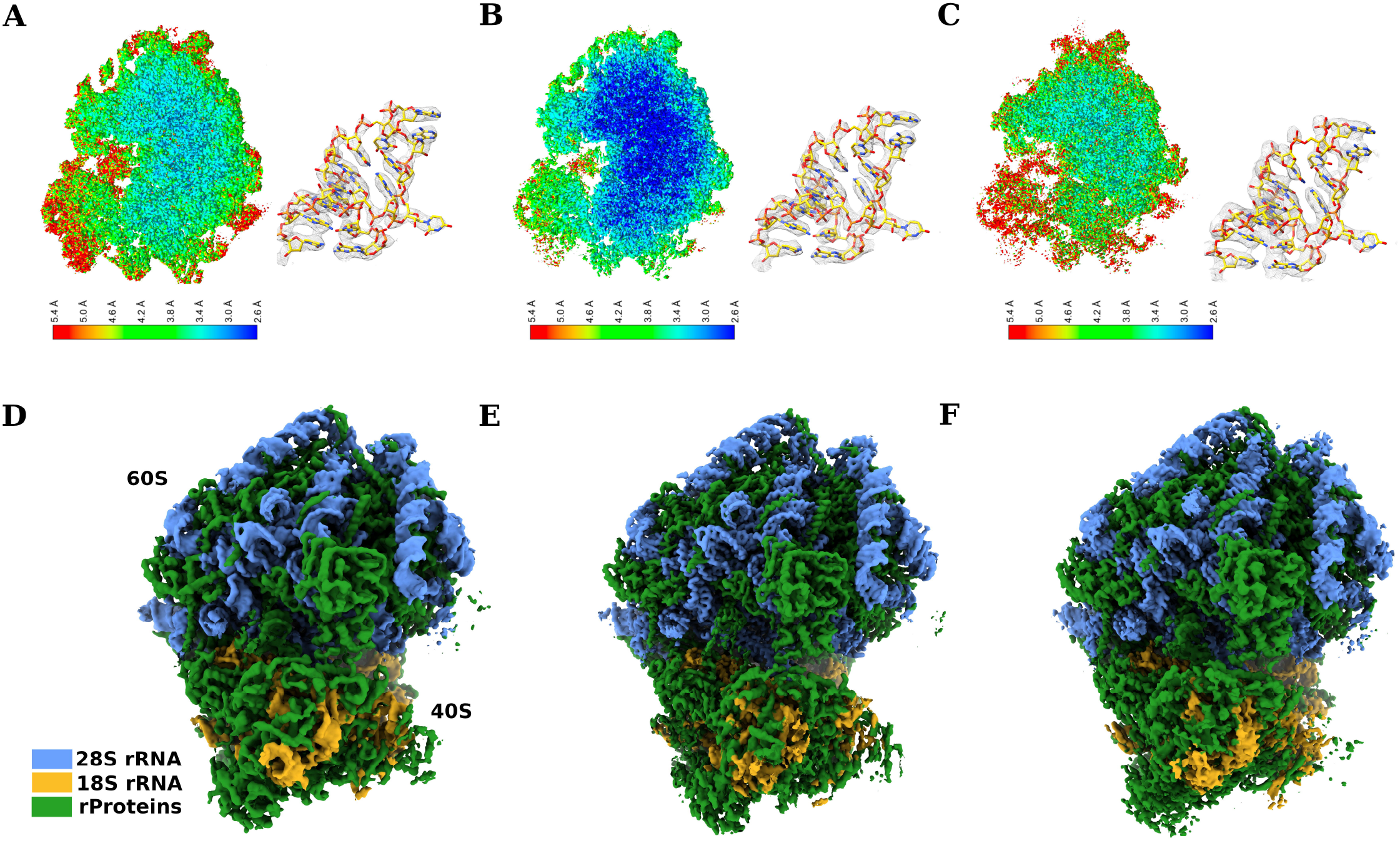
(A) Local resolution map of the NMR ribosome, highlighting a selected portion of 28S rRNA residues 2385 to 2388 with corresponding density. (B) Same as (A) for tuco-tuco ribosome. (C) Same as (A) for guinea pig ribosome. (D) Overall map of the NMR ribosome, colored based on identities of rRNA and ribosomal proteins. (E) Overall map of tuco-tuco ribosome, same coloring scheme. (F) Overall map of guinea pig ribosome, same coloring scheme.

We identified at least one ribosome class with clear density for endogenous mRNA in the mRNA cleft for all three species: naked mole rat, tuco tuco, and guinea pig. This allowed us to directly model native mRNA segments into the cryo-EM maps, providing structural snapshots of ribosomes actively engaged in translation. To our knowledge, these are the first high-resolution structures of natively purified, actively translating ribosomes from any mammalian species. Prior structures from native sources usually captured idle or post-termination states without mRNA in the decoding site.

Interestingly, both the NMR and tuco-tuco ribosomes display the ribosome in an eEF2-bound state without indications of any mRNA, which may have been lost during purification. Although we could not locate any bound GTP or GDP on eEF2, the entire protein could be modeled with confidence, including the essential domain IV required for effective translocation (Figure 2). Consistent with observations in human ribosomes, our density maps for both NMR and tuco-tuco ribosomes exhibit clear map coverage for the mammalian-specific insertion within the eEF2 G′ domain (not shown). While the density for the diphthamide moiety of eEF2 was not well resolved in our structures, we were able to identify the conserved His715 residue unambiguously in both the NMR and tuco-tuco eEF2 models (Figures 2D, E, F).

**Figure 2:**
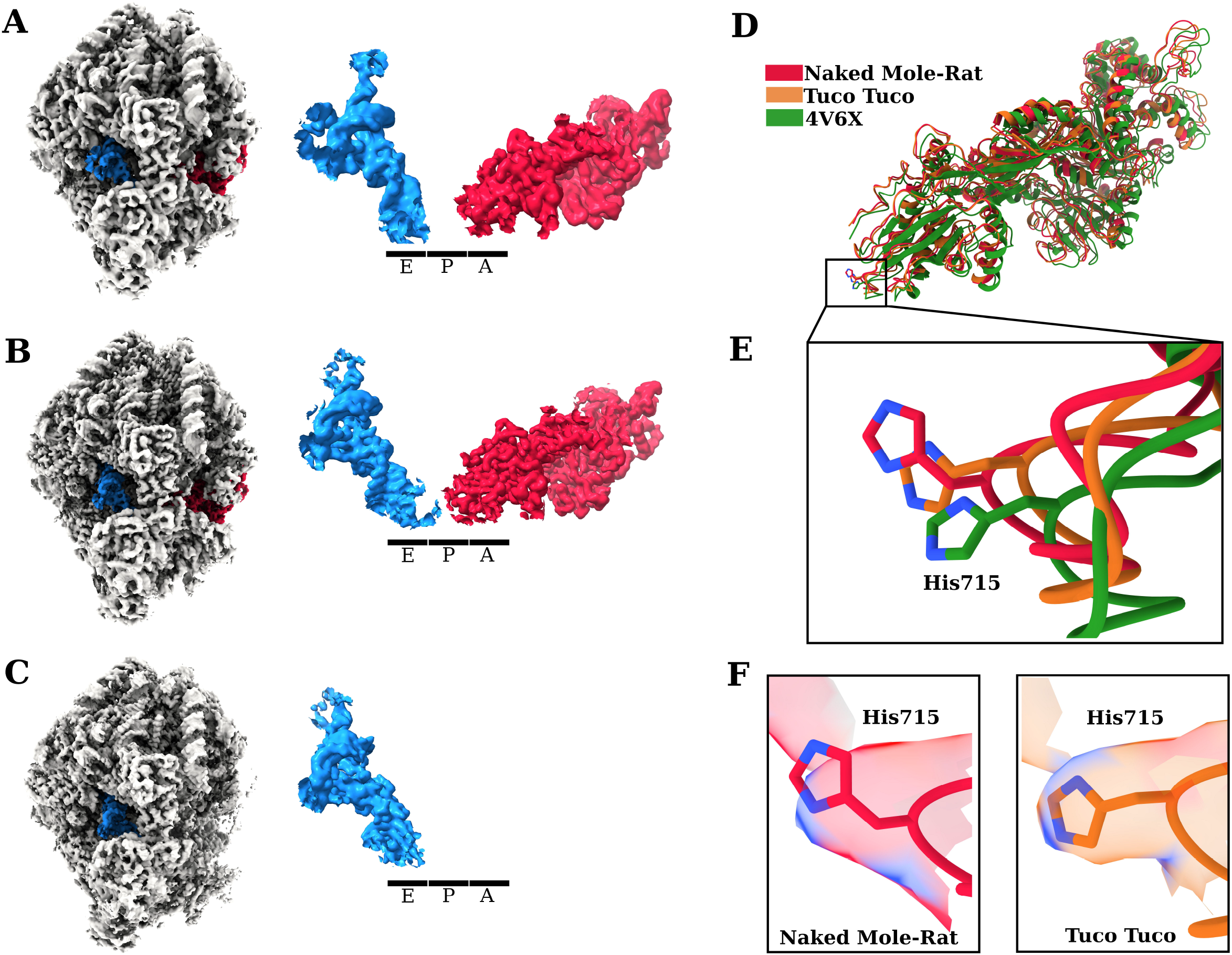
(A) Density map of NMR ribosome bound with P/E-tRNA and eEF2, with zoomed-in view showing the interaction between tRNA and eEF2. (B) Same as (A) for tuco-tuco ribosome. (C) Same as (A) for guinea pig ribosome, except no eEF2 density is observed in this structure. (D) Structural comparison of eEF2 from NMR and tuco-tuco ribosomes with eEF2 from PDB 4V6X (human). (E) Zoomed-in view of His715, the residue that undergoes diphthamide modification for NMR, Tuco-tuco, and human (4V6X). (F) In all structures, density is only observed for the histidine moiety, with no clear density for the modified diphthamide group. This may be due to the absence of mRNA in our structures, causing the modified moiety to be flexible and disordered.

A comparative analysis of the position of eEF2 His715 within the 80S ribosome revealed species-specific differences. When aligned with the human ribosome structure (PDB ID: 4V6X), the His715 residue in our NMR-eEF2 complex was found to be ~3.8 Å closer to the P site, while in the tuco-tuco-eEF2 complex it was ~2.8 Å closer (Figure 2E). Due to the absence of eEF2 in our guinea pig maps, we were unable to include this species in the comparison.

The NMR ribosome closely resembles other eukaryotic ribosomes, most closely that of rabbit (Bhatt et al., 2021; De et al., 2024). Its ribosomal proteins share over 95% sequence identity with both rabbit and guinea pig ribosomes (Table 2). Some sequence variations are observed in a few large subunit (LSU) proteins, namely uL4, eL6, eL14, and eL29. However, these variations do not result in significant structural changes. Overall, the ribosome structure is highly conserved, with minor differences primarily at the terminal ends of these proteins, which exhibited weak cryo-EM density due to mobility and could not be fully modeled.

Interestingly, these LSU proteins interact with rRNA expansion segments (ES7L, ES9L, ES15L, and ES39L), which exhibit considerable variation across species. Sequence variations in the terminal ends of these proteins are likely coupled with subtle changes in rRNA in a way that preserves the functionality of the ribosome involving the dynamic interplay between rRNA and ribosomal proteins.

In the tuco-tuco ribosome, unique adaptations of the ribosomal components are also evident, particularly in the LSU proteins uL4, eL6, eL14, and eL29. This species shows an extended ES15L compared to the NMR.

Additionally, we closely examined the appearances of Helix 69 (H69) and the GTPase-associated center (GAC) across the three ribosome structures determined in this study, alongside previously published human and rabbit ribosome structures. The analysis was performed by aligning both the 28S rRNA and the entire 80S ribosome structures separately, followed by calculation of the root-mean-square deviation (RMSD) between each pair. H69 has been implicated in substrate selection within the ribosome (Ortiz-Meoz and Green, 2011); however, our analysis revealed no significant differences in its conformation among the three rodent ribosomes. In contrast, the GAC, comprising H43 and H44 and connected to the main body of the large subunit via H42, is highly dynamic and only poorly resolved in our cryo-EM density maps, making detailed comparisons impossible.

While H69—known for its role in substrate selection (Ortiz-Meoz and Green, 2011)—remained conserved, the GAC (comprising H43, H44, and connected via H42) exhibited high dynamism and was poorly resolved in our cryo-EM maps, precluding detailed comparisons.

### 3. 28S rRNA fragmentation and ES15L in naked mole-rat and tuco-tuco ribosomes

The splicing of the rRNA transcripts for the large ribosomal subunit (LSU) 28S of NMR and tuco-tuco leads to the fragmentation of the latter into two distinct RNA segments. We adopt the designation LSU-α and LSU-β, as previously defined for similar rRNA fragments in *Trypanosoma* by Hashem et al. (2013).

In the NMR, there are two specific cleavage sites within the divergent region 6 (D6) of the 28S rRNA transcript, leading to the excision of a 263-nucleotide fragment (Azpurua et al., 2013). This cleaving process splits the 28S rRNA into LSU-α and LSU-β fragments of unequal sizes. In contrast, the tuco-tuco rodent (*Ctenomys* genus) has a single cleavage site within a unique 106-base pair insertion in the D6 region, thereby forming its fragments LSU-α and LSU-β (Melen et al., 1999; Azpurua et al., 2013). In both species, the two fragments are peripherally located; LSU-α is primarily on the solvent side of the 60S subunit, and LSU-β at the 40S-60S interface.

Our density map of the NMR ribosome shows high-resolution details of the fragmentation site near ES15L, which caps H45. This region is stabilized by base-stacking interactions involving LSU-α and ribosomal protein eL28. In contrast, the tuco-tuco ribosome retains an extended ES15L, which projects outward from the 60S subunit’s solvent-side surface (Figure 3).

**Figure 3:**
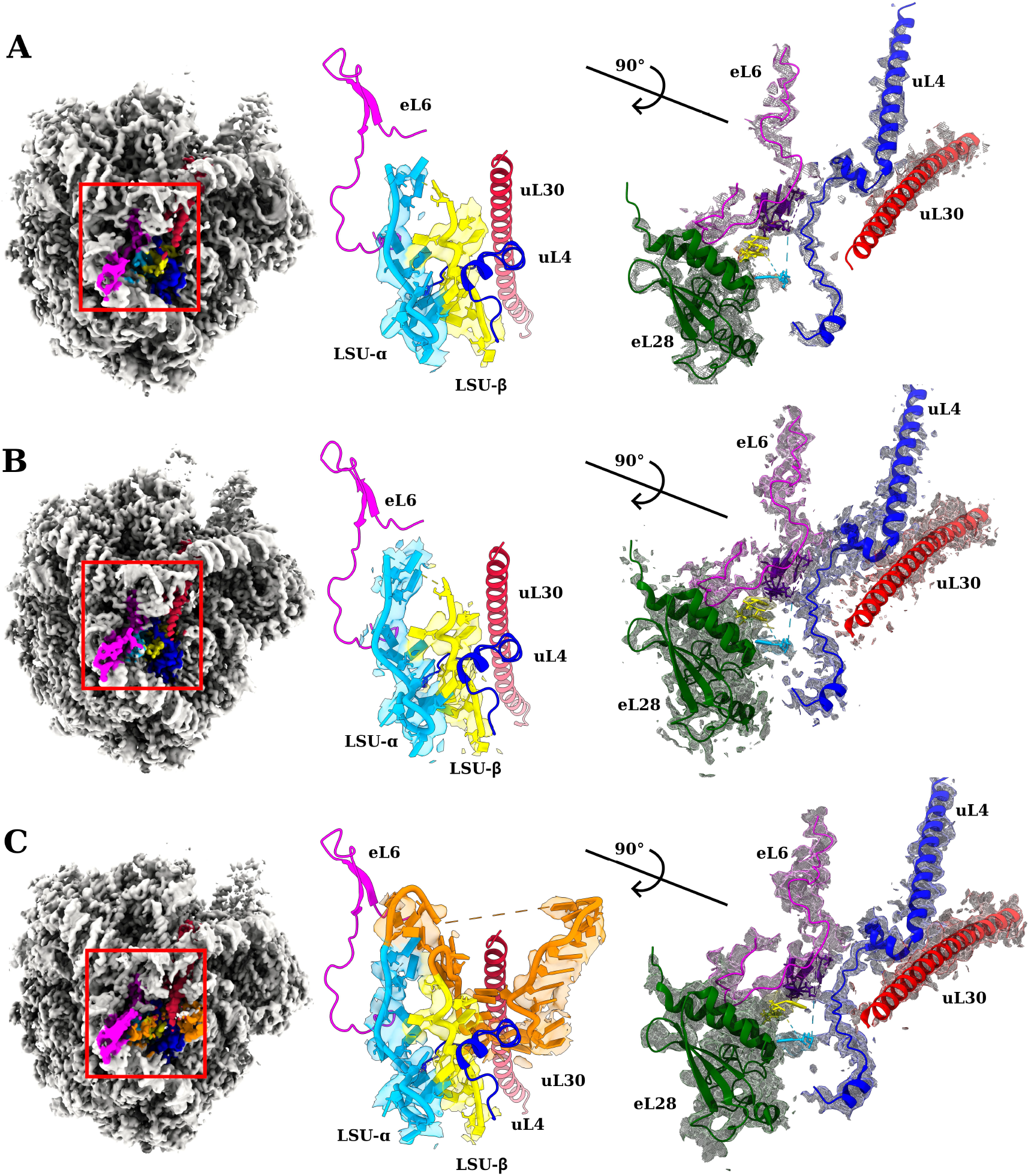
(A) Architecture of the 28S rRNA fragmentation site in the NMR ribosome, shown with an overview (first column) and two detailed zoomed-in views. The second column zooms into the red-boxed region, displaying only the transparent cryo-EM map surface to highlight the modeled RNA. The view is from the top of the 60S subunit, with density shown only for the LSU-α and LSU-β fragments. The map coverage ends precisely at the fragmentation junctions, aligning with the predicted cleavage sites for this species. The third column shows a 90° rotated view focused on surrounding ribosomal proteins, whose positions and map coverage appear nearly identical to those in mammalian ribosomes without fragmented rRNA. (B) Same as (A) for the Tuco-tuco ribosome. (C) Same as (A), shown for the guinea pig ribosome. Unlike the NMR and tuco-tuco, the guinea pig ribosome does not exhibit rRNA fragmentation, and the corresponding region (colored in orange) is clearly resolved in our structure. In the rightmost panel, the rRNA segment that is excised in NMR and Tuco-tuco is not shown to avoid obstruction and to visualize the surrounding ribosomal proteins better. These proteins occupy similar positions and are resolved similarly in our maps across all three species, indicating that rRNA fragmentation in NMR and Tuco-tuco does not lead to major overall structural changes in the ribosome.

In human ribosomes, the terminal loop of H30 within ES9L forms a continuous stretch of base pairs with an internal region of ES15L, creating a hybrid ES9L–ES15L helix that anchors the base of the human-specific ES15L extension firmly to the ribosomal surface (Anger et al., 2013). In contrast, such interactions are absent in the NMR and tuco-tuco ribosomes due to ES15L fragmentation. The ES9L-ES15L hybrid helix appears to be maintained in the guinea pig ribosome since we observe a helically shaped mass of density in the unsharpened map; however, this part of the structure could not be modeled due to the low resolution in this peripheral region. The functions of ES15L, or the ES9L–ES15L hybrid helix, are not fully understood. To date, the only experimentally demonstrated, validated role of ES15L is in 60S ribosomal subunit biogenesis (Ramesh and Woolford et al., 2016).

We compared the fragmented 28S rRNA of the NMR and tuco-tuco ribosomes with the canonical, non-fragmented 28S rRNA in guinea pig. Aside from the missing nucleotides, minor shifts in adjacent helices, and slight adjustments in expansion segments in the fragmented 28S rRNA, no major structural differences are observed. Given the A-site finger’s (ASF) known role in maintaining translational fidelity (O’Connor et al., 1997; Takeo et al., 2006) and its physical connection to the fragmented helix of ESL15L, we closely examined the ASF structures in all three ribosome maps. Our analysis revealed that the ASFs are of similar length, structure, position, and conformation across all three species (not shown).

### 4. Comparison of rRNA Expansion Segments Across Species

Our comparative analysis of ribosomal expansion segments in the NMR, tuco-tuco, guinea pig, and human ribosomes show both conserved features and structural differences (Figure 4). Expansion segment ES7L is present in all species but is slightly shorter in the tuco-tuco ribosome map compared to guinea pig and human ribosomes. Similarly, ES27L is shorter in tuco-tuco than in the NMR and guinea pig. Due to the lack of a reliable sequence for these rRNAs, structural models were based on visual interpretation of cryo-EM maps, following a strategy similar to that of Bhatt et al. (2021).

**Figure 4:**
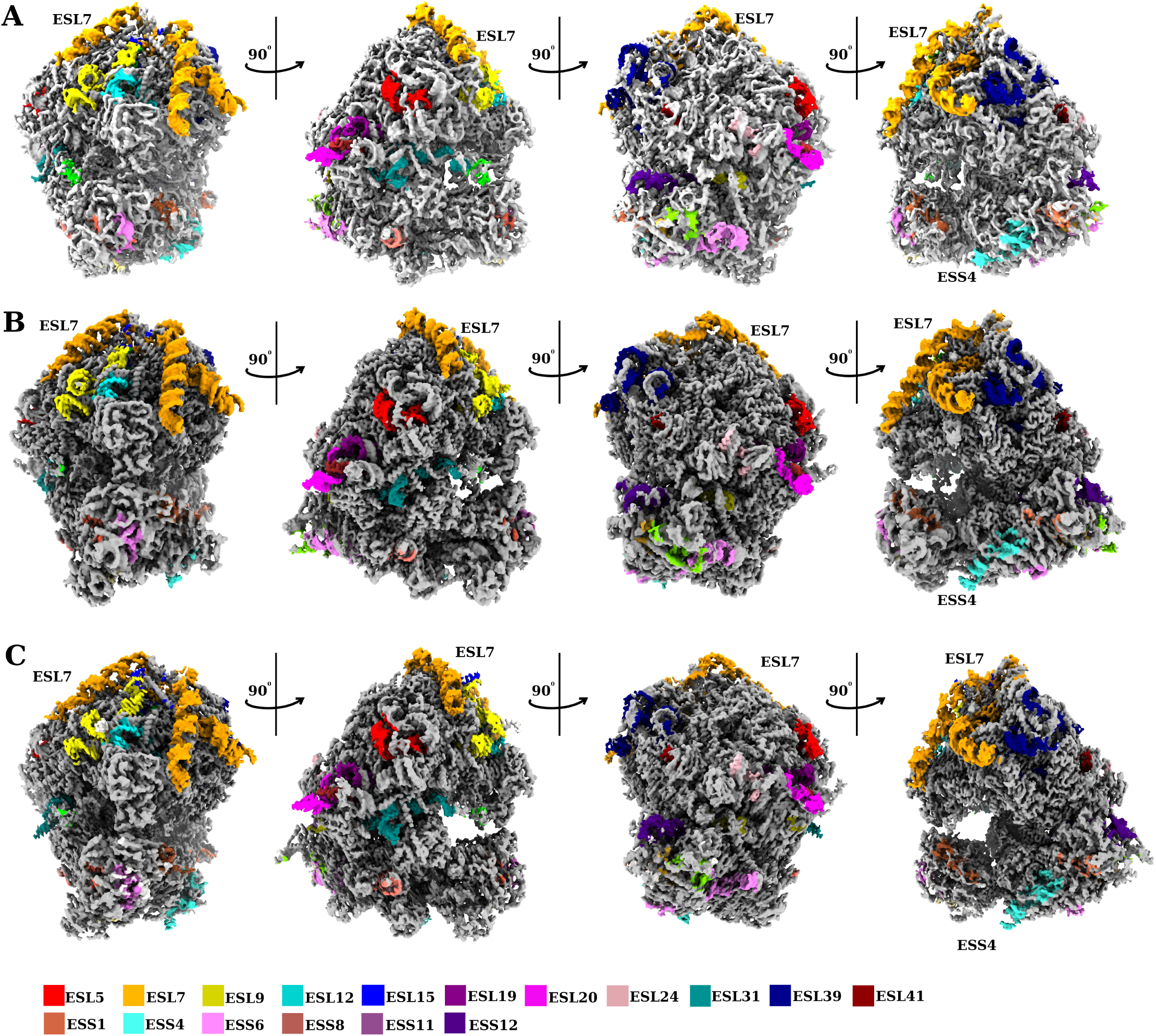
(A) Expansion segments in NMR ribosome. (B) Expansion segments in tuco-tuco ribosome. (C) Expansion segments in guinea pig ribosome.

In both NMR and tuco-tuco, all expansion segments show up with weak density, particularly ES8L, indicating high mobility, i.e., positional heterogeneity unresolved by classification. Similarly, ES10L and ES12L appear as mobile regions in rodent ribosomes.

The structure of ES15L, which interacts with ES27L, appears conserved across the three species. However, the tuco-tuco ribosome exhibits shorter ES15L and ES27L regions compared to the NMR. At very low RMSD or threshold, ES15L in the NMR ribosome remains visible up to position 2156, with the returning helix appearing from 2218. In contrast, in the tuco-tuco ribosome, ES15L is visible only up to position 2102, with the helix reappearing starting at 2246. A similar pattern, though with less stark differences, is observed for ES27L, where the NMR ribosome maintains coverage up to position 2952 and resumes at 3242, while in the tuco-tuco ribosome, these positions shift to 2946 and 3246, respectively. In comparison, ES15L and ES27L in the guinea pig and NMR ribosomes appear more or less similar in length and structure. In human and *Drosophila* ribosomes, ES31L-A and ES27L form additional inter-subunit bridges, which were also resolved at high resolution in the three rodent structures analyzed. These extended helices are absent in yeast and other lower eukaryotes.

### 5. Modified tRNA residues

In the structural analysis of the ribosomes, we observed distinct modifications and structural features associated with the tRNA across species. tRNA modifications such as wybutosine at position 37 play a crucial role in stabilizing the anticodon loop, ensuring proper codon recognition, and preventing frameshifting during translation (Rak et al., 2021). These modifications also contribute to maintaining translational accuracy and decoding efficiency by enhancing base-stacking interactions and restricting the flexibility of the anticodon (Schweizer et al., 2017; Xu et al., 2017). In class 2 of the NMR ribosome map, we identified a mass of density corresponding to the YYG37 modification, located near the 18S ribosomal RNA at nucleotide 1249U with a proximity of 2.9 Å. In contrast, no clear density is observed for the Ψ39 modification, and the anticodon region appears unstructured. In the tuco-tuco ribosome, the YYG37 modification is also discernable, although residues 33–38 are unstructured in the map. For guinea pig class 1, the ribosome displays the YYG37 modification interacting with the 18S ribosomal RNA at a closer distance of 2.8 Å, along with additional density coverage for the anticodon loop. In addition, the guinea pig class-2 map shows the E-site tRNA in a slightly different conformation compared to our maps for NMR and tuco-tuco.

## Discussion

Unlike the guinea pig and other mammals studied to date, the NMR and tuco-tuco possess fragmented rRNA, and this distinct feature prompts intriguing questions regarding the evolutionary and functional implications of rRNA fragmentation in mammals. Particularly, we are looking for properties of the NMR’s rRNA structure that could explain the organism’s exceptional biological traits—high translational fidelity, longevity and stress resistance. Its high fidelity is also shared with tuco-tuco.

The presence of endogenous mRNA in our reconstructions underscores the physiological relevance of our findings, opening new avenues to study translation in its native regulatory and compositional context. Our high-resolution structural analyses of NMR and tuco-tuco ribosome-eEF2 complexes reveal detailed aspects of the mammalian eEF2-ribosome interaction. Notably, our work, like the human ribosome study (Anger et al., 2013), utilized native purification after puromycin treatment. Observed differences in eEF2 positioning, particularly the proximity of His715 to the P site compared to the human structure, could suggest potential species-specific variations in the eEF2-ribosome interaction landscape across mammals.

Does the fragmentation of rRNA lead to higher fidelity by changing the dynamics of the ribosome? To look for possible clues supporting this hypothesis, we examined the dynamics of specific ribosomal elements in both NMR and tuco-tuco, as reflected by the way they appear in the density maps, in comparison with the way they appear in guinea pig. The elements we singled out were the central protuberance, A-site finger, helix 69 (H69), and the GTPase-associated center, all part of 28S rRNA. Dynamics of these elements accompany key processes, such as mRNA-tRNA translocation, during the elongation phase of translation. However, neither our structural results nor inferences on dynamics from weak densities give evidence that the enhanced fidelity observed in NMR and tuco-tuco can be attributed to rRNA fragmentation. It is still possible, however, that it might arise from subtle allosteric effects not observable in our analysis.

Among the most striking findings in our study is the fragmentation pattern of the 28S rRNA in both NMR and tuco-tuco. In NMR, the 28S rRNA is composed of two pieces of unequal sizes, termed LSU-α and LSU-β, resulting from the excision of a 263-nucleotide segment from the full-length sequence at two cleavage sites. In contrast, tuco-tuco’s 28S rRNA transcript undergoes only a single cleavage without the excision of any rRNA segment. Despite these structural modifications, both species evidently retain the core functional architecture of the ribosome. One point of possible interest is that the fragmentation of ES15L in the NMR prevents its interaction with ES9L. This prevents the formation of the ES9L–ES15L hybrid helix observed in human and guinea pig ribosomes. The absence of this helix in the NMR and tuco-tuco suggests that the stabilizing role of ES9L is unnecessary when ES15L is truncated—i.e., when its length and flexibility are reduced. The function of ES15L remains incompletely understood; it has been implicated in 60S ribosomal subunit biogenesis (Ramesh and Woolford et al., 2016) and, more speculatively, proposed to participate in mRNA recruitment (i.e., translational regulation) (Parker et al., 2018). If ES15L contributes to mRNA recruitment or interactions with regulatory factors, its truncation may influence which transcripts are preferentially translated—or alternatively—may reflect a loss of this regulatory capacity altogether. Supporting a potential regulatory role, Rodríguez-Algarra et al. (2025) identified a dense cluster of intragenomic variants within human ES15L, that associate with diverse body size-related traits such as height, weight, and cholesterol levels. These findings suggest that ES15L variation may not be neutral but could contribute to phenotypic diversity, possibly through structural or regulatory impacts on ribosome function. However, such possibilities remain highly speculative given the limited experimental evidence beyond its established role in biogenesis.

The A-site finger, whose dynamics have been linked to the maintenance of translational fidelity (O’Connor et al., 1997; Komoda et al., 2006), does not show any structural differences among the NMR, tuco-tuco, and guinea pig ribosomes. Additionally, there is no evidence in the cryo-EM maps suggesting differences in the dynamic features of this region.

Expansion segments (ESs) in ribosomal RNA are important for ribosome assembly, translation accuracy, and mRNA selection (Fujii et al., 2019; Biesiada et al., 2022; Rauscher et al., 2024; Rauscher and Polacek, 2024). Specific ESs, such as ES7S and ES27L, enhance decoding fidelity and influence nascent peptide modification, highlighting their diverse functional roles (Shankar et al., 2020; Lindahl 2024). Thus, high similarity in the structure and interactions of ESs indicates that translational regulation in two species is closely matched.

A key observation of our study is the near-identical structure of rRNA ESs in guinea pig and human ribosomes. Another key finding is the conservation of the expansion-segment cluster formed by ES7L, ES9L, ES10L, and ES15L between human and guinea pig ribosomes. Although most ESs in human ribosomes are significantly longer than those in guinea pig ribosomes, these length differences do not substantially alter the conserved interactions within the cluster.

This high similarity between human and guinea pig ribosomes contrasts with the marked differences observed in tuco-tuco and NMR ribosomes. In tuco-tuco, ES15L is visible only up to residue 2123 and resumes at 2252, while in NMR, it is even more limited, with visibility up to 2124 and resuming at 2255. Additionally, ES27L exhibits disordered helices and is poorly resolved in the tuco-tuco maps, with only a short stretch (residues 2894–2901) visible. In contrast, in NMR and guinea pig, ES27L is clearly visible from residues 2894–2915, and the map coverage of remainder of the helix is also present, albeit at lower resolution.

Our maps from the NMR and tuco-tuco indicate mobility in the peripheral regions of ES10L, ES15L, as well as the C-terminal portion of ribosomal protein eL29, which are all poorly resolved in the density maps. Specifically, the fragmentation of ES15L in the NMR ribosome prevents its interaction with helix 30 (H30) from ES9L, a feature that distinguishes it from both the tuco-tuco and guinea pig ribosomes.

Future research, particularly involving ribosomes bound to both cognate and near-cognate tRNAs, could provide additional insights into whether rRNA fragmentation contributes to the accuracy of tRNA selection or error correction. Further studies could also investigate how this fragmentation affects the ribosome’s interaction with various translational factors, or with quality control mechanisms like ribosome-associated quality control (RQC), in maintaining translational accuracy. Considering the NMR’s unique adaptations, such as robust proteostasis, existence of efficient DNA repair mechanisms, and resistance to oxidative stress, it is plausible that rRNA fragmentation might also play a role in preserving proteome integrity, particularly in the context of the species’ long lifespan and stress resilience.

The importance of studying ribosomal variability among model organisms extends to the realm of drug development, where even minor structural differences in ribosomes can influence interactions with small-molecule drugs. While ribosomes are generally conserved across species, even slight variations in structure or dynamics may result in different drug responses, potentially affecting the accuracy of preclinical studies. By understanding these subtle differences, researchers can improve the precision of drug testing, enhance predictions for human therapeutic responses, and ultimately refine preclinical research methodologies.

In conclusion, our study shows remarkable structural diversity in mammalian ribosomes. It tried to address the significance of rRNA fragmentation, particularly its possible link to the unusual resilience and longevity of two species. The diversity of ribosomal architecture across the rodent species investigated highlights the need for continued investigation. Future research focusing on the functional implications of ribosomal variability, particularly regarding proteome maintenance and stress resilience, will be critical for understanding the evolutionary and biological rationale of rRNA fragmentation in the NMR and tuco-tuco. This line of inquiry could yield valuable insights into the mechanisms that support the unique physiological traits of these species and may offer broader implications for translational research, drug development, and our understanding of ribosome evolution.

## Author contributions

The project was conceived by A.S., V.G. and J.F.. Z.K. provided cell cultures and protein harvests. C.G. prepared cryo-EM grids and collected data with supervision and assistance from Z.L. Data processing, structure determination, atomic model building and visualization were initially carried out by C.G. with assistance by Z.L. and M.N., then revised and refined by S.D. with assistance from C.G.. Detailed analysis, inter-species comparisons of ribosome structures and analysis of conformational mobility were done by C.G., S.D. and J.F.. rRNA sequence comparisons were performed jointly by C.G., S.D., M.N. and S.M.. Final figures were prepared by C.G. and S.D.. Project administration and supervision of the biochemistry experiments were provided by A.S. and V.G., for the cryo-EM team by J.F.. The manuscript was written by C.G., S.D., V.G. and J.F., with input from A.S., Z.K., and S.M.. All authors reviewed and approved the final manuscript.

## Acknowledgements

This work was supported by grants from the U.S. National Institute on Aging (P01AG047200 to A.S. and V.G.) and the U.S. National Institutes of Health (R35GM139453 to J.F.). Cryo-EM data were collected at Columbia University’s Cryo-EM Core and the HHMI Janelia Cryo-EM Facility. RNA sequencing was performed at the JP Sulzberger Columbia Genome Center. We thank Bob Grassucci for assistance with data collection and Xiaohan Hu (University of Bristol) for help with model curation and deposition.

## Methods

### 1. Cell culture and ribosome purification

Naked mole-rat (NMR) fibroblasts were grown at 32 °C (in vivo body temperature) 5% CO2, 3% O2 on treated polystyrene culture dishes in eagle’s minimum essential medium (EMEM) media (ATCC) supplemented with 15% FBS, nonessential amino acids, sodium pyruvate, 100 U/mL penicillin, and 100 µg/mL streptomycin (Gibco). Cells were centrifuged and the supernatant removed. Pellets were stored at −80 °C. Tuco-tuco fibroblast cells were grown in similar conditions with temperature adjusted to 37 °C.

Fibroblast cell pellets were thawed and resuspended in a lysis buffer containing 50 mM Tris (pH 7.4), 300 mM potassium acetate, 7 mM magnesium acetate, 380 mM sucrose, 2 mM DTT, 0.14% (v/v) Triton X-100, and a Roche protease inhibitor tablet. After 20-minute incubation on ice, cells underwent two freeze-thaw cycles for complete lysis. The lysate was clarified by centrifugation at 11,700 x g until no pellet remained.

Ribosomes in the supernatant were pelleted through a 1 M sucrose cushion in 20 mM HEPES (pH 7.4), 300 mM potassium acetate, 7 mM magnesium acetate, 2 mM DTT, and 20% glycerol, by centrifugation at 40,000 rpm for 16 hours at 4°C in a 70.1 Ti fixed-angle rotor. The ribosome-enriched pellet was resuspended in 200 µl of the same buffer, clarified by centrifugation, treated with 1mM puromycin for 30 minutes on ice to improve homogeneity of the sample. The sample was then loaded onto a two-layer sucrose cushion (20% and 40% w/v sucrose in 20 mM HEPES, pH 7.4, 100 mM potassium acetate, 10 mM magnesium acetate, and 2 mM DTT). The ribosome-enriched pellet was resuspended in 200 µl of the same buffer above but without sucrose.

For the guinea pig sample, a sucrose gradient (10-50%) replaced the two-layer cushion, yielding a distinct ribosomal peak. The gradient was prepared with a BioComp Gradient Master and centrifuged for 15 hours at 19,000 rpm at 4°C. Fractions containing 80S ribosomes were collected, dialyzed to remove sucrose, and concentrated to ~600 nM.

### 2. Cryo-EM Sample Preparation

Purified ribosomes (3 µl) were applied to copper/holey carbon grids (carbon-coated Quantifoil R2/2 grids), coated with an additional thin carbon layer and glow-discharged. Grids were blotted for 3 seconds at 4°C and 100% humidity before plunge-freezing in liquid ethane cooled to −180°C. For the guinea pig sample, Quantifoil R 1.2/1.3 300 mesh Au holey carbon grids were used.

### 3. Imaging and Data Collection

NMR and tuco-tuco ribosomes were imaged on a Titan Krios microscope with a Falcon 2 camera (HHMI Janelia Research Campus) at a magnification of 133,970x, yielding a pixel size of 1.045 Å. Each image, collected in integrating mode, consisted of 25 frames with a total dose of 40 e□/Å^2^. The guinea pig sample was imaged using an FEI Tecani F30 Polara microscope equipped with a Gatan K2 Summit camera.

### 4. Image Processing

Micrographs were inspected for contamination and beam shifts prior to movie processing and particle picking. Image processing involved frame alignment, particle picking, 2D and 3D - classifications, particle polishing, and CTF estimation.

3D classification, auto-refinement, and post-processing were carried out using RELION (Scheres, 2012), yielding resolutions of 3.2 Å for NMR ribosomes and 3.0 Å for the large subunit. Approximately 400,000 particles from the NMR dataset underwent 3D classification to remove impurities, followed by multiple classification rounds to separate ribosome conformations and refine homogeneity.

Three distinct ribosome conformations were identified and refined to high resolution (3.2 Å for the 80S ribosome and 3.0 Å for the large subunit). Further refinement involved 2D and 3D classification, 3D auto-refinement with particle polishing, MotionCor2 (Zheng et al., 2016), and per-particle CTF refinement. Focused 3D refinement achieved a 3.0 Å resolution for the large subunit.

Modeling for NMR, guinea pig, and tuco-tuco ribosomes was performed in Coot (Emsley and Cowtan, 2004) and refined in Phenix (Liebschner et al., 2019). Due to the lack of curated rRNA and ribosomal protein sequences for these species, model building required alternative strategies. For the naked mole-rat, we used partial RACE sequencing data mapped onto a rabbit ribosome model as a reference. For tuco-tuco, which lacks a reference genome or curated transcriptome, we relied on our RNA-seq data and used the rabbit model as a structural and sequence guide. We were able to extract the tuco-tuco rRNA sequences with confidence and incorporate them into the structural models. However, the RNA-seq data yielded fragmented ribosomal protein sequences; therefore, we retained the rabbit r-protein sequences for model building. In the case of the guinea pig, we directly used rabbit ribosome sequences due to their high similarity and availability. These comparative approaches allowed us to build accurate and well-refined models despite the limited availability of species-specific sequence information.

### 5. Workflow for Ribosomal RNA and Protein Sequence Extraction from RNA-Seq Data

Total RNA was extracted from tuco-tuco fibroblast cells, as previously described (Azpurua et al., 2013), and submitted to the JP Sulzberger Columbia Genome Center. RNA quality was assessed using a BioAnalyzer (Agilent Technologies; Instrument ID: DE72901373, Firmware: C.01.069, Type: G2939A), and the sample yielded an RNA integrity number (RIN) of 7.7. Either this total RNA was used directly for library preparation, or poly-A selection was performed to enrich for mRNA. Libraries were constructed using Illumina TrueSeq chemistry and sequenced using Illumina NovaSeq 6000. Base calling was performed by the Columbia Genome Center using Illumina’s Real-Time Analysis (RTA) software.

Our workflow for extracting ribosomal RNA (rRNA) and ribosomal protein sequences from RNA-Seq data integrates standard bioinformatic tools with customized steps to ensure efficient and accurate identification of these crucial translational components. The initial phase involves preparing the reference genome and the necessary rRNA databases. Specifically, the *Oryctolagus cuniculus* (rabbit) genome and its annotation were downloaded from Ensembl. The genome was indexed for rapid sequence access using samtools faidx and for efficient alignment with the STAR aligner. Concurrently, comprehensive rRNA databases were obtained from the SortMeRNA repository, encompassing sequences from SILVA, Rfam, and other sources. These diverse rRNA sequences were combined into a single indexed FASTA file, optimized for rapid alignment of sequencing reads against known rRNA.

The subsequent stage focused on processing the raw RNA-Seq data and aligning it to the prepared reference genome. Raw FASTQ files underwent quality assessment using FastQC to evaluate read quality metrics. Adapter sequences and low-quality bases were then removed using Trim Galore, ensuring the removal of technical artifacts before alignment. The cleaned reads were then aligned to the indexed rabbit genome using STAR, a splice-aware aligner designed for RNA-Seq data. Post-alignment quality control was performed using tools like RSeQC and dupRadar to assess read distribution and duplication levels, resulting in high-quality aligned BAM files suitable for downstream analyses.

To specifically isolate reads originating from ribosomal RNA, the workflow employed SortMeRNA. This high-performance tool aligns the quality-filtered FASTQ files against the precompiled and indexed [all_rRNA.fasta] database. SortMeRNA utilizes a seed-based alignment algorithm to efficiently and sensitively identify rRNA sequences within the dataset. The output of this step is a clear separation of the sequencing reads into an rRNA-aligned fraction and a non-rRNA fraction. This separation is critical for downstream transcriptomic analyses that often focus on the non-rRNA portion of the transcriptome, while the extracted rRNA reads are available for further characterization.

In contrast to rRNA identification, ribosomal protein-coding sequences were extracted directly from the complete set of RNA-Seq reads without prior rRNA removal. This process begins with the alignment of the processed sequencing reads to the rabbit reference genome using STAR, as described earlier. Following alignment, read quantification is performed using featureCounts to assign reads to specific genomic features. A curated list of gene identifiers belonging to the RPS (small subunit), RPL (large subunit) families and other ribosomal protein coding genes, was utilized in conjunction with the genome annotation file to isolate ribosomal protein-coding genes. By intersecting the aligned reads with the genomic regions annotated as ribosomal protein genes, we were able to extract the reads specifically mapping to these sequences, enabling the quantification of individual ribosomal protein gene expression levels.

## FIGURE LEGENDS

**Table 1:** Cryo-EM and Refinement Statistics

**Supplementary Figure 1:** CryoEM image processing workflow

**Supplementary Figure 2:** CryoEM FSC curves

**Supplementary Figure 3:** RNAseq workflow

