## Supplementary figures and images for "Structures of Naked Mole-Rat, Tuco-Tuco, and Guinea Pig Ribosomes–Is rRNA Fragmentation Linked to Translational Fidelity?"

### Supplemental Fig1

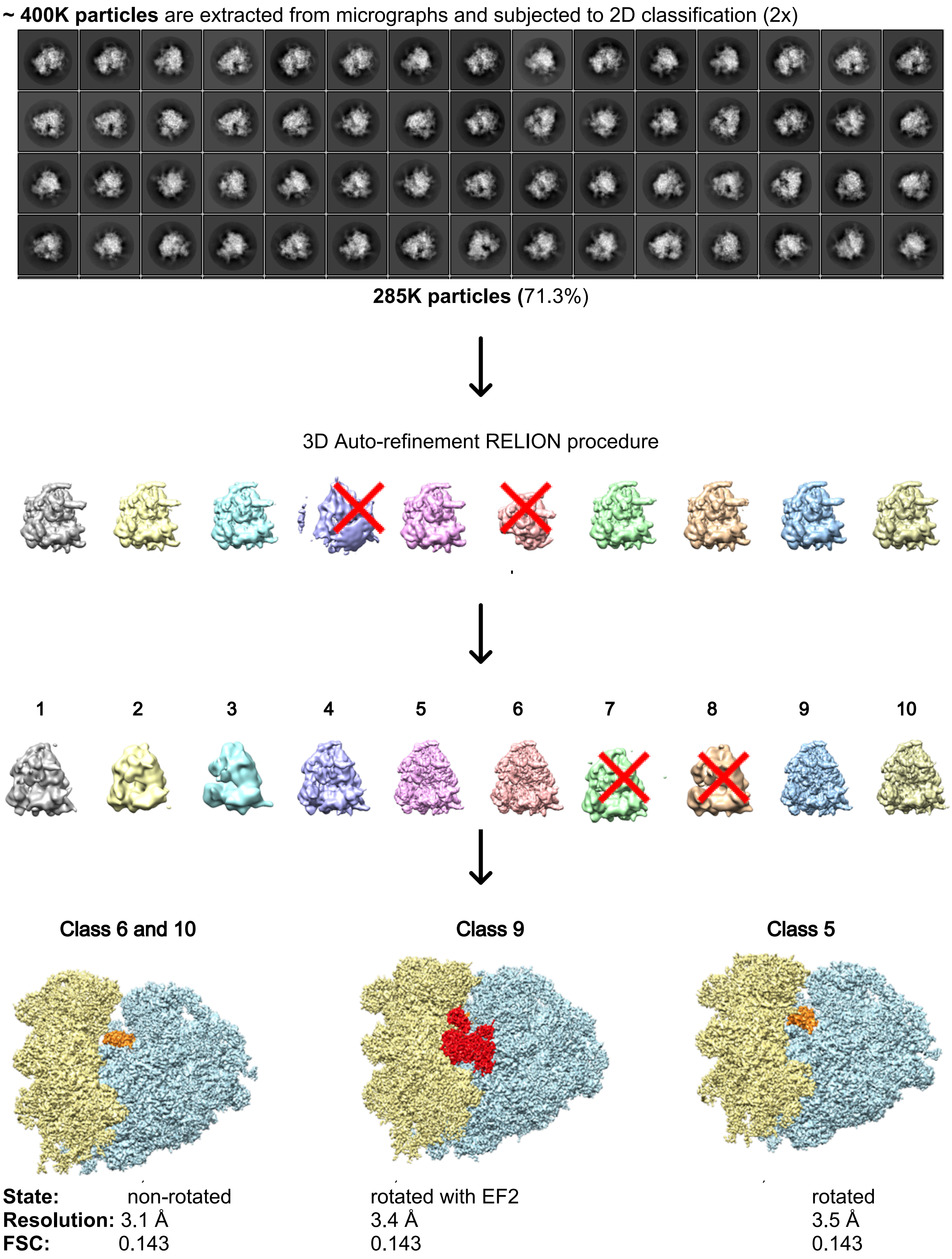

### Supplemental Fig2

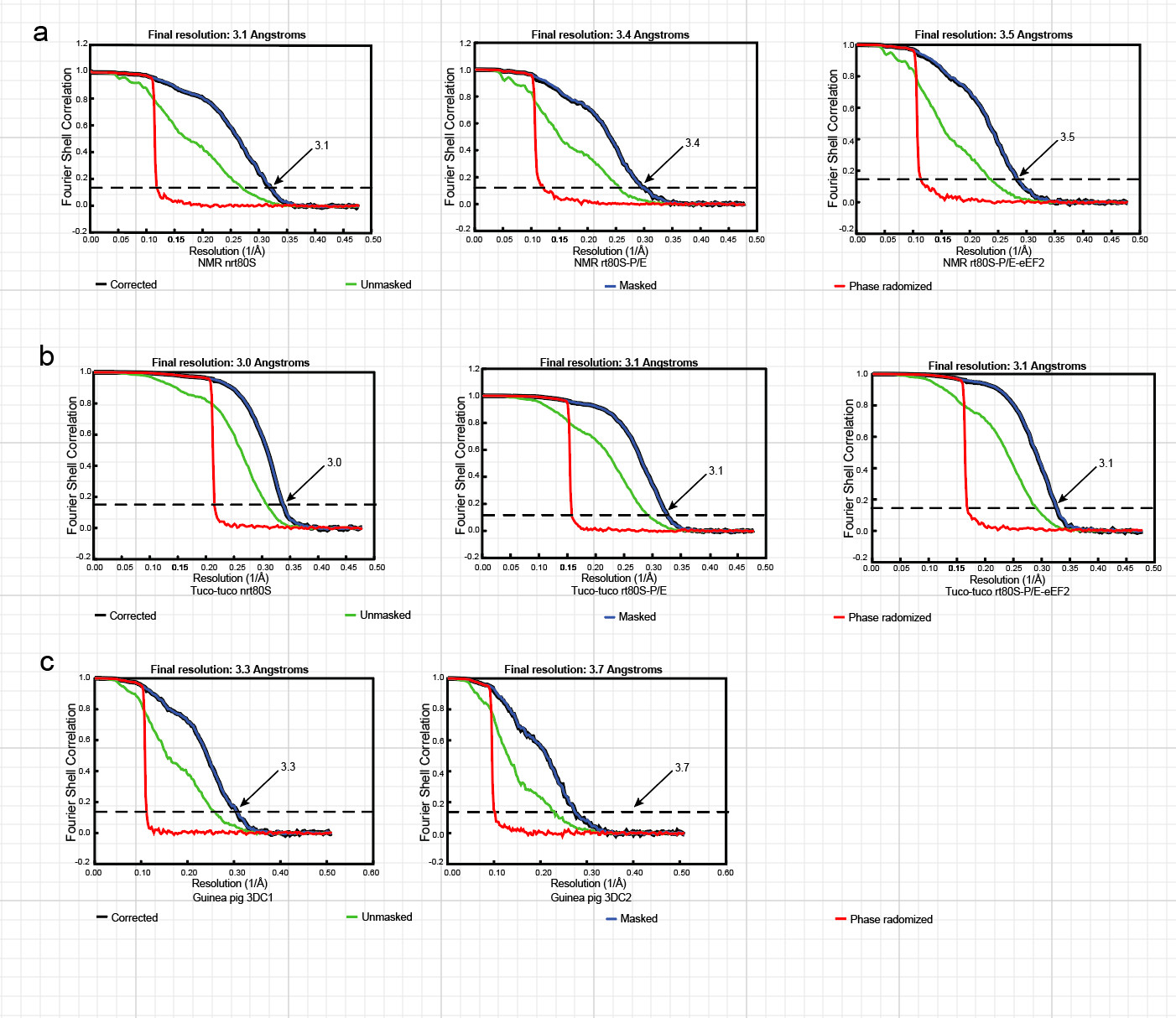

### Supplemental Fig3

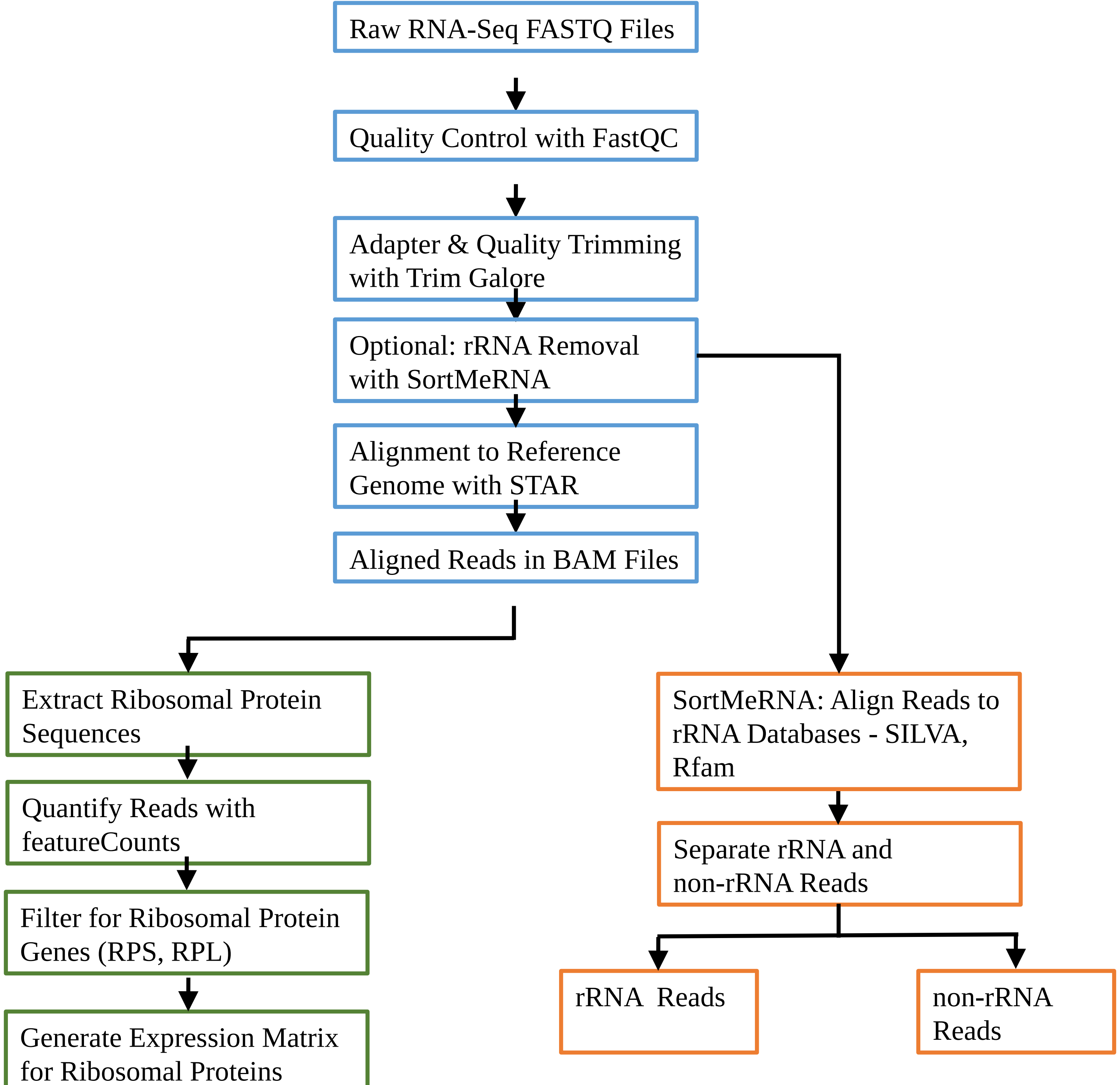
