## Supplementary material for "Structures of Naked Mole-Rat, Tuco-Tuco, and Guinea Pig Ribosomes–Is rRNA Fragmentation Linked to Translational Fidelity?": Table 1

**Cryo-EM data collection, refinement and validation statistics**

|  | NMR non-rotated | NMR with  eEF2 | TT  rotated | TT with  eEF2 | GP non-rotated |
| --- | --- | --- | --- | --- | --- |
| **Data collection and processing** |  |  |  |  |  |
| Scope and Camera  Magnification | Krios,  Falcon2  133,970 | Krios,  Falcon2 133,970 | Krios,  Falcon2 133,970 | Krios,  Falcon2  133,970 | Polara F30, K2  39,000 |
| Voltage (kV) | 300 | 300 | 300 | 300 | 300 |
| Electron exposure (e–/Å^2^) | 42 | 42 | 23 | 23 | 71 |
| Defocus range (μm) | 3.0 to -1.0 | -3.0 to -1.0 | -3.0 to -1.0 | -3.0 to -1.0 | -3.0 to -1.0 |
| Pixel size (Å) | 1.045 | 1.045 | 1.045 | 1.045 | 0.95 |
| Symmetry imposed |  |  |  |  |  |
| Initial particle images (no.) | C1  400000 | C1 400000 | C1 400000 | C1 400000 | C1 400000 |
| Final particle images (no.) |  |  |  |  |  |
| Map resolution (Å)  FSC threshold | 3.1 0.143 | 3.4 0.143 | 3.3 0.143 | 4.1 0.143 | 4.8 0.143 |
| Map resolution range (Å) |  |  |  |  |  |
| **Refinement** |  |  |  |  |  |
| Initial model used (PDB code) | 5LZS/ 7O7Y | 5LZS/ 7O7Y | 5LZS/ 7O7Y | 5LZS/ 7O7Y | 5LZS/ 7O7Y |
| Map sharpening *B* factor (Å^2^) | – | – | – | – | – |
| Model composition  Non-hydrogen atoms  Protein residues  Nucleotide residues  Ligands | 217120  12153 5748  Zn:8 Mg:276 | 217120  11924 5652  Zn:8 Mg:275 | 225771  12554 5678  Zn:8 Mg:275 | 220934  12153 5748  Zn:8 Mg:275 | 225771  12777 5744  Zn:8 Mg:275 |
| *B* factors (Å^2^)  Protein  Ligand |  |  |  |  |  |
| R.m.s. deviations  Bond lengths (Å)  Bond angles (°) | 0.003 0.692 | 0.007 1.110 | 0.005 0.924 | 0.004 0.754 | 0.005 0.669 |
| Validation  MolProbity score  Clashscore  Poor rotamers (%) | 1.91 3.61 3.23 | 1.99 5.82 3.08 | 2.07 4.80 3.30 | 1.78 3.28 3.43 | 2.51 3.79 2.04 |
| Ramachandran plot  Favored (%)  Allowed (%)  Disallowed (%) | 94.50 5.10 0.40 | 95.72 4.19 0.09 | 93.42 6.34 0.24 | 96.28 3.68 0.04 | 96.02 3.92 0.06 |
